# Detecting aberrant splicing events in short-read RNA-seq with SAMI, an UMI-aware Nextflow pipeline

**DOI:** 10.1101/2025.03.10.642333

**Authors:** Sylvain Mareschal, Valentin Wucher, Sarah Huet, Camille Léonce, Kaddour Chabane, Sandrine Hayette, Pierre-Paul Bringuier, Stéphane Pinson, Marc Barritault, Claire Bardel

## Abstract

**Summary:** Despite RNA-sequencing solutions have existed for 15 years, successfully replacing micro-arrays for gene expression profiling, their potential to analyse splicing is still to be achieved in clinical settings. Powerful tools were developed to quantify known isoforms, to compare sample groups or to identify outliers in large datasets, but most fail to address the most common situation in routine diagnostics: identifying non-recurring events in low-dimension data. To fill this gap, we developed SAMI (Splicing Analysis with Molecular Indexes), an UMI-aware pipeline focusing on splicing events which differ from provided annotation. The ability of SAMI and concurrent software to detect intragenic splicing aberrations and gene fusions was assessed, both on real data from a commercial control sample and simulated data generated with ASimulatoR.

**Availability and Implementation:** Nextflow pipeline and Singularity container recipe freely available under GPL 3 licence at https://github.com/HCL-HUBL/SAMI

## 1 Introduction

Splicing analysis has been addressed by numerous tools in research contexts, where biological replicates from a limited set of predefined conditions can be gathered (rMATS [1], SUPPA2 [2], MAJIQ [3], …). Clinical diagnostic requires however to conclude on individual and uncharacterized samples, an issue that tools like SPOT [4], LeafCutterMD [5] or FRASER 2 [6] tried to address by looking for outliers in large cohorts of samples. These tools rely on statistical and/or denoising models which require high dimension data (both in terms of screened genes and “normal” samples to compare to), conditions which are hard to meet in facilities running disease-specific sequencing panels.

SAMI was developed to address such a need, i.e. maximizing sensitivity in small panels where samples may be scarce or likely to harbor recurring events. As Splice-Launcher [7] has been used for such a purpose in french facilities, it served as a comparison point for SAMI performances. SAMI (Splicing Analysis with Molecular Indexes) was moreover designed to take full benefits from currently available sequencing panel kits, handling Unique Molecular Indexes (UMIs) and stranded sequencing.

## 2 Implementation

SAMI is an integral Singularity-contained Nextflow [8] pipeline, processing raw FASTQ files directly from the sequencer (Supp Figure 1). It covers usual RNA-seq processing steps such as adapter trimming with cutadapt [9] and two-pass alignment to the genome with STAR [10], and provides multiple quality checks gathered with MultiQC [11]. These quality controls include FastQC [12], Picard’s CollectRnaSeqMetrics [13], featureCounts [14] and several custom controls.

SAMI takes full benefit from the two-pass alignment performed by STAR, which has been shown to improve the accuracy of splicing discovery [15]. Splicing gaps are collected by STAR from each sample separately during the first pass, and injected into the reference genome to provide consistent gaps among samples during the second pass. SAMI further refines this gap collection by shifting novel junctions toward annotated splicing sites when multiple alignments are possible, enhancing significantly the accuracy over alternative 5’/3’ splicing aberrations.

This two-pass approach also provides an opportunity to efficiently handle Unique Molecular Indexes (UMIs), which are now part of several commercial RNA-seq kits and allow for a more accurate deduplication than sequence-based approaches [16]. In SAMI, mapping co-ordinates from the first pass are used by fgbio [17] to generate consensus reads from UMI clusters, which are aligned during the second pass. In the absence of UMIs, SAMI will run a classical two-pass STAR mapping.

Splicing analysis is performed by a collection of scripts integrated in SAMI. Evidence of splicing is collected from BAM files by an HTSlib-based [18] C program, along with read pair orientation. Junctions are then annotated using Rgb [19] and the provided reference transcriptome, and classified according to the status of the two splicing sites involved :

- Annotated: both sites are annotated, and at least one annotated transcript holds it;
- Plausible: both sites are annotated, but no transcript exists with this junction;
- Anchored: only one site is annotated;
- Unknown: none of the sites is annotated.

This approach allows SAMI to detect both intra-gene splicing and gene fusion events, and particular care was taken on STAR parameters to achieve high sensitivity for both applications.

To filter candidate events, SAMI provides two PSI (Percent Spliced In) values for each junction in each sample, reflecting how frequently it occurs among all splicing observed at the two involved splicing sites. *PSI* values provided here are restricted to a single *junction* in the context of a single splicing *site*, as opposed to tools like rMATS [1] which try to integrate multiple evidence sources into pre-defined splicing events. The support of a splice at a specific site *reads*_*site*_ is the amount of split reads evidencing a mapping gap starting or ending at this site.

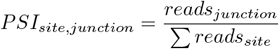

SAMI further extends this PSI-based approach by considering the absence of splice at a site (as observed in intron retention or exon elongation) as an alternative during PSI computation. This is achieved by measuring the sequencing depth 3 bases inside the putative intron, i.e. after or before the considered splicing site, for all annotated and discovered splicing sites. These alternatives add two more classes for junctions returned by SAMI :

- No-splice: absence of splice at an annotated splice site;
- Trivial: absence of splice at a de novo splice site (inside an annotated intron or exon);

As final products, SAMI provides both filtered and unfiltered junction tables with extensive annotation, and graphical representations of each gene in each sample which evidenced at least one filtered junction of interest (Supp Figure 2).

## 3 Performances

### 3.1 Seraseq^®^ commercial samples

A benchmark on the Seraseq sample has been performed using a panel of 21 genes, which intersects two splicing events (skipping of exon 14 in *MET* and of exons 2 to 7 in *EGFR*) and 15 detectable gene fusions of clinical interest (Supp Table 1). Seven samples were processed in clinical conditions with a regular RNA quantity of 10 ng and six samples with an increasing RNA quantity: 6.25 ng, 10 ng, 12.5 ng, 25 ng, and two samples with 50 ng.

While both tools have a high mean number of false positives without filtering, 731 and 371 for SAMI and Splice-Launcher respectively, this number is greatly reduced by minimal filtering (41 and 56 respectively, Figure 1a). All expected events are found in all samples without filtering, but one event with low support is missed once an intermediate (SAMI) or a stringent (SpliceLauncher) filter is used. This can be explained by the different definitions of the thresholds depending on the internal thresholds of each tools. Moreover, the amount of false positives can be increased by the annotation used to predict the splicing events. Indeed, if a splicing event is physiological but not annotated, it will be considered as a false positive.

**Figure 1:**
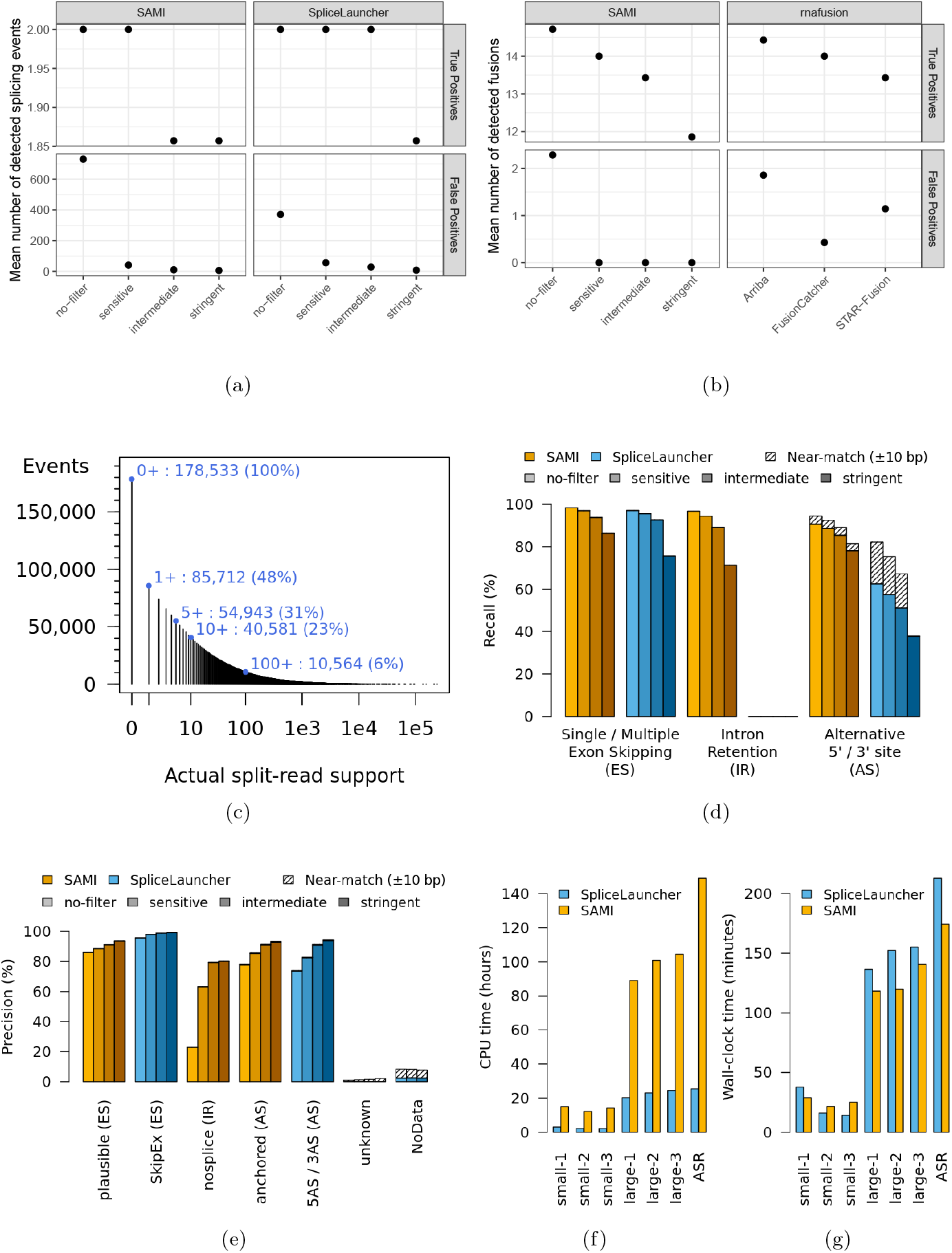
SAMI’s performances, compared to SpliceLauncher and rnafusion. **(a)** Average of True (TP, top panel) and False (FP, bottom panel) positive splicing events detected on the seven SeraSeq samples containing two expected splice events (skips of MET exon 14 and EGFR exons 2 to 7) for SAMI (left panel) and SpliceLauncher (right panel) using four thresholds: no-filter, sensitive, intermediate, and stringent. **(b)** Average of True (TP, top panel) and False (FP, bottom panel) positive fusion events detected on the seven SeraSeq samples containing 15 expected fusion events for SAMI (left panel) and tools from rnafusion (right panel). For SAMI, four thresholds have been used: no-filter, sensitive, intermediate, and stringent. For rnafusion, the predictions of three tools are shown: Arriba, FusionCatcher, and STAR-fusion. **(c)** Amount of splicing events generated by ASimulatoR supported by at least the number of split-reads in X. **(d)** Proportion of splicing alterations created by ASimulatoR which were detected by SAMI and SpliceLauncher, for each ASimulatoR event class and for various parameters of the two tools. Events found by a tool with slightly incorrect genomic coordinates (10 bp difference or less) were counted separately as near-matches. **(e)** Proportion of splicing alterations reported by SAMI and SpliceLauncher which were actually generated by ASimulatoR, split by event classes reported by each tool and for various parameters. **(f)** CPU and wall-clock computation time spent running SAMI or SpliceLauncher on 3 real-life runs of two RNA-seq panels of different sizes and on data generated by ASimulatoR.

Regarding gene fusion detection, SAMI was compared to the nf-core pipeline [20] “rnafusion” version 3.0.2, which includes Arriba [21], FusionCatcher [22] and STAR-fusion [23], using fastp trim and the default parameters. While all methods have similar performances in terms of true positives, SAMI didn’t detect any false positive event even with a sensitive threshold (Figure 1b). On increasing quantity of Seraseq RNA, SAMI reached its recall plateau at 10 ng without a filter and at 25 ng with the most permissive filter, while the other methods did so at 10 ng (Supp Figure 3).

### 3.2 Simulated data

To provide a broader overview of SAMI’s and Splice-Launcher’s capabilities regarding splicing, RNA-seq data with known aberrations was generated using ASimulatoR [24]. FASTQ files were produced to simulate 16 million pairs of 100 bp reads per sample from MANE Select transcripts of the RefSeq GRCh38 transcriptome. 30 samples were produced through 10 simulations, each simulation expressing at various levels 8,000 genes harboring each a single event of exon skipping, multiple-exon skipping, intron retention, alternative 5’ or 3’ splicing site (12.5% chance each) or no event (37.5% chance). The whole cohort was analyzed by SAMI and SpliceLauncher with default parameters and full RefSeq GRCh38 annotation.

As ASimulatoR synthetically expresses genes at a wide range of levels, 92,821 (52.0%) of the 178,533 simulated events were not supported by any generated split read and were thus impossible to detect (Figure 1c). Focusing on 40,581 events supported by at least 10 simulated reads, SAMI’s recall with “intermediate” filtering was 93.8%, 89.1% and 85.4% for exon skips (ES), intron retentions (IR) and alternative 5’/3’ sites (AS) respectively (Figure 1d), with corresponding precision values of 91.1%, 79.4% and 90.9% (Figure 1e). Similar ES values were obtained with SpliceLauncher (92.6% recall and 98.9% precision), but recall was significantly lower on AS (51.0%) and IR are out of the tool’s scope. Up to 19.7% of AS events were not reported accurately by SpliceLauncher (shifted by 10 bp or less and improperly classified) but still present in the results, theoretically raising recall to 67.2% with “intermediate” filtering.

As opposed to SAMI, SpliceLauncher works with a single transcript definition per gene, which fits well this simulation setting. In results presented here (Figures 1d and 1e) the transcripts used for simulation were indicated to SpliceLauncher, but performances were significantly lower when they were not (Supp Figures 4 and 5).

### 3.3 Computation time

SAMI and SpliceLauncher ran on a dedicated computing node exposing 900 Go of RAM and 110 computing threads from two Intel^®^ Xeon^®^ Platinum 8352M CPUs to a SLURM instance. Computing time was assessed for 3 runs of a small MiSeq panel (16×0.8 million read pairs from 21 loci), 3 runs of a larger NextSeq panel (16×10 million read pairs from 165 genes) and a single run of ASimulatoR synthetic data (30×16 million read pairs from 8,000 genes).

SpliceLauncher proved to be 4 to 6 times lighter than SAMI in terms of CPU time (Figure 1f), but efficient parallelization in SAMI resulted in similar wall-clock durations (Figure 1g). The higher computing intensivity of SAMI could be explained mainly by quality checks not performed by SpliceLauncher, more complex handling of UMIs and different parameters for STAR alignment (Supp Table 2).

## 4 Conclusion

SAMI is a powerful tool for the detection of aberrant splicing events in disease-specific RNA-seq panels. Compared to SpliceLauncher, it offers equivalent performances in the detection of exon-skipping events, significantly better ones on alternative 5’/3’ splicing sites and moreover detects intron retentions and gene fusions. Its implementation as an heavily-parallelized Singularity-contained Nextflow pipeline, and optional compatibility with UMIs and stranded RNA-seq kits, make it a portable and modern solution to handle RNA-seq data in a clinical context.

## 5 Author contributions

**Conceptualization:** SM, VW, SHu, PPB, SP, MB, CB; **Formal analysis / Methodology / Software / Original draft:** SM, VW; **Funding acquisition:** SHu, MB, CB; **Resources:** SHu, SHa, SP, MB, CB; **Project administration / Supervision:** MB, CB; **Investigation:** CL, KC, SHa; **Review & editing:** all authors.

