## Supplemental Methods, Tables and Figures for "Detecting aberrant splicing events in short-read RNA-seq with SAMI, an UMI-aware Nextflow pipeline"

March 10, 2025

### 1 Supplemental Methods

#### 1.1 SAMI

All analyzes were performed with SAMI version 2.1.0, using GCA_000001405.15_GRCh38_full_analysis_set reference genome and RefSeq annotation from 2024-08-23. The 4 profiles described in the article correspond to the following launch parameters :

```
no-filter:    --min.I=1  --min.PSI=0
sensitive:    --min.I=3  --min.PSI=0.01
intermediate: --min.I=5  --min.PSI=0.05
stringent:    --min.I=10 --min.PSI=0.1
```

Annotated events, i.e. "annotated" and "trivial" classes, have been ignored during analyzes.

#### 1.2 SpliceLauncher

All analyzes were performed using a fork of https://github.com/LBGC-CFB/SpliceLauncher [1] from commit `1ca5dc72` (2022-07-01), adding minor adaptations to local computing infrastructure. Singularity recipe and annotation were built following the provided README on 2022-09-20. Unfortunately issues with latest versions of SpliceLauncher were not solved at the time of publication and prevented the benchmark to be performed with more recent versions.

As SpliceLauncher focuses on a single transcript for each gene, which is the last in alphabetical order for the version used here, a `--transcriptList` file was provided with the MANE transcript for each gene and the `--removeOther` flag was not used.

For real-life data including Unique Molecular Indexes (UMIs), a deduplication step with CReaK from the Agilent Genomics NextGen Toolkit (AGeNT) version 3.1.2 was performed between separate calls to SpliceLauncher for alignment and counting, with the following arguments : `--consensus-mode="SINGLE" --remove-dup-mode -f --MBC-mismatch=1 --min-avg-MBC-qual=20 --min-avg-read-qual=20 -F -MS`.

The 4 profiles described in the article correspond to the following post-processing filters :

```
no-filter:    sample count >= 1
sensitive:    sample count >= 3
intermediate: sample count >= 5  & sample expr >= 0
stringent:    sample count >= 10 & sample expr >= 1
```

Annotated events, i.e. "Physio" types, have been ignored during analyzes.

#### 1.3 ASimulatoR

RNA-seq data with known splicing events was generated with ASimulatoR [2] version 1.0.0 running in R version 4.0.4. Transcripts were extracted from `GCA_000001405.15_GRCh38_full_analysis_set` RefSeq annotation from 2024-08-23, excluding all scaffolds except chromosomes 1 to 21, X and Y. ASimulatoR pools exons from all annotated transcripts at the gene level, using chimeric transcripts never observed in the annotation as "templates" and thus introducing unexpected splicing aberrations. To bypass this issue, only one transcript (the one selected by MANE) was used for each gene. `simulate_alternative_splicing()` was called 10 times with the following arguments :

```
event_probs = c(es=0.125, mes=0.125, ir=0.125, a3=0.125, a5=0.125)
multi_events_per_exon = FALSE
max_genes = 8000
num_reps = rep(1, 3)
meanmodel = TRUE
readlen = 100
paired = TRUE
seq_depth = 10e6
adapter_contamination = TRUE
pcr_rate = 0.3
distr = "empirical"
error_model = "illumina5"
bias = "none"
strand_specific = TRUE
gzip = TRUE
```

As event annotation and counts provided by ASimulatoR proved to be unreliable, split-reads supporting exon junctions were recounted from transcript coordinates provided by ASimulatoR in read names. Template and variable transcripts described in `splicing_variants.gtf` were systematically compared to identify novel junctions induced by exon skips (ES), multiple consecutive exon skips (MES), alternative 5' (a5) and 3' (a3) splicing sites. Similarly, the two expected "no-splice" events induced by each intron retention (IR) were inferred from these transcripts. Recall of SAMI ans SpliceLauncher was computed as the ability to retrieve these novel junctions, looking at genomic coordinates rather than transcript annotation.

#### 1.4 Seraseq® commercial sample

18-plex Seraseq Fusion RNA Mix v4 reference standard sample was purchased from LGC Seracare (Milford, USA) and sequenced multiple times with the "small" RNA-seq panels described below. The two splicing events and 16 gene fusions expected in this control samples are described in Supp Table 2.

#### 1.5 "Small" RNA-seq panel

RNA was extracted using the Maxwell® RSC RNA FFPE Kit (Promega, Reference AS1440) and the Maxwell® RSC Instrument (Promega, Catalog Number AS4500) from Formalin-Fixed, Paraffin-Embedded (FFPE) samples. According to manufacturer instructions, RNA libraries were prepared using the KAPA RNA HyperPrep Kit in combination with the KAPA Universal UMI Adapter with a sample input of 10 ng. The workflow involved pre-PCR for 18 cycles, followed by target enrichment using the KAPA HyperPETE LC Fusion Panel, an 18 kb capture target panel that includes 17 lung cancer fusion genes and 4 housekeeping genes as internal controls. Libraries were captured using the KAPA HyperPETE Reagent Kit and sequenced on an Illumina Miseq® System.

### 2 Supplemental Figures

#### 2.1 Supp Figure 1: Overview of SAMI's workflow

The main processes of SAMI are illustrated with blue boxes, sample-specific inputs and outputs with dotted lines and run-level ones with solid lines. Processes generating QC data aggregated in the final MultiQC reports are marked with a magnifying glass icon.

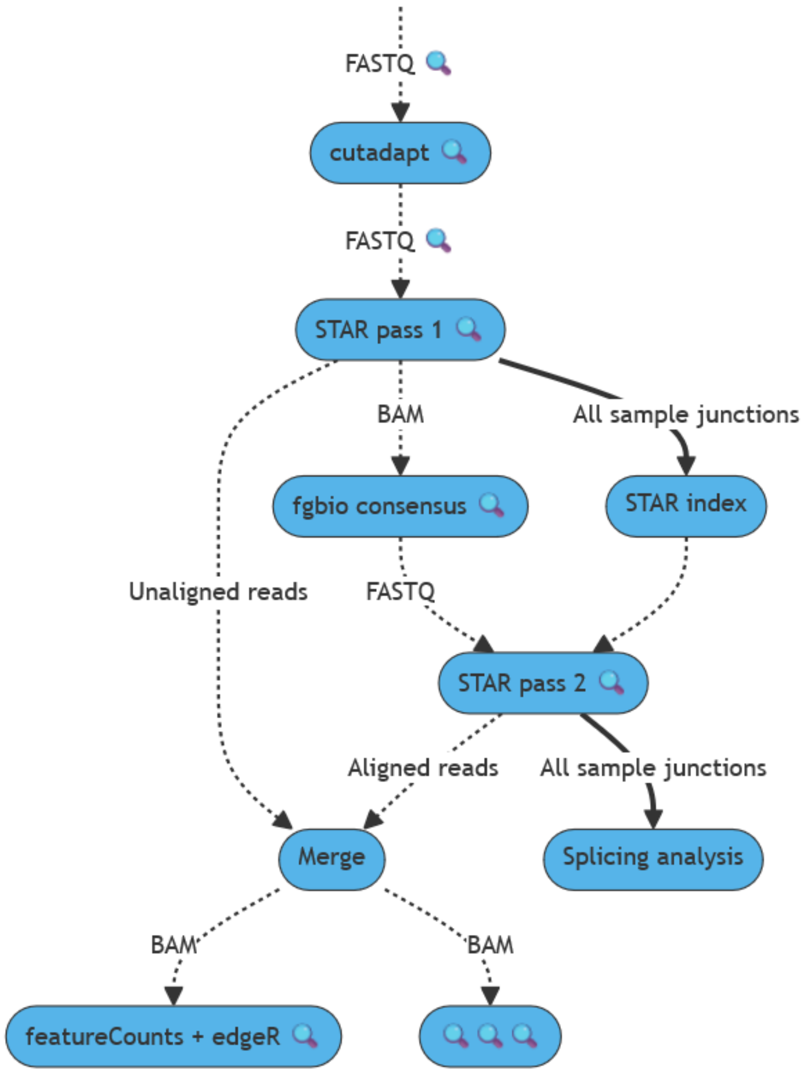

#### 2.2 Supp Figure 2: Example of plots generated by SAMI

SAMI produces one plot for each gene (here ENG) and each sample harboring at least one event passing filters. The three transcripts described in the annotation for the considered gene are presented in the middle, with square boxes representing exons (numbered here in reverse order as ENG is transcribed from the reverse genomic strand). Exons are shaded according to the average sequencing depth observed in the sample, from lowest observed value in white to deepest coverage in black. Blue bridges on the top side illustrate the amount of split reads supporting annotated junctions, while colored bridges at the bottom represent aberrant splicing observed in the sample.

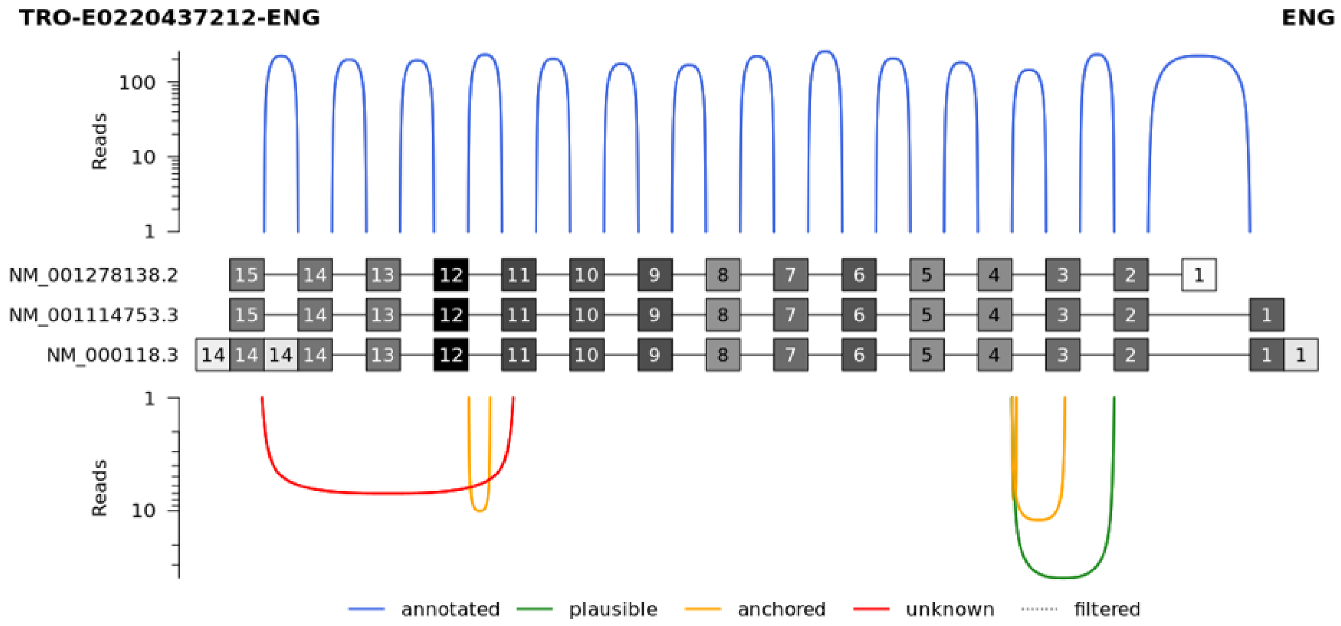

#### 2.3 Supp Figure 3: True and False positive fusion events

Amount of True (plain lines) and False (dashed lines) positive fusion events detected on the six SeraSeq samples with an increasing RNA quantity, 6.25 ng, 10 ng, 12.5 ng, 25 ng, and two samples with 50 ng, containing 15 expected fusion events for SAMI (top panels) and tools from rnafusion (bottom panel). For the 50 ng quantity, the mean value of the two samples is represented. For SAMI, four thresholds have been used (from left to right): no-filter, sensitive, intermediate, and stringent. For rnafusion, the predictions of three tools are shown (from left to right): Arriba, FusionCatcher, and STAR-fusion.

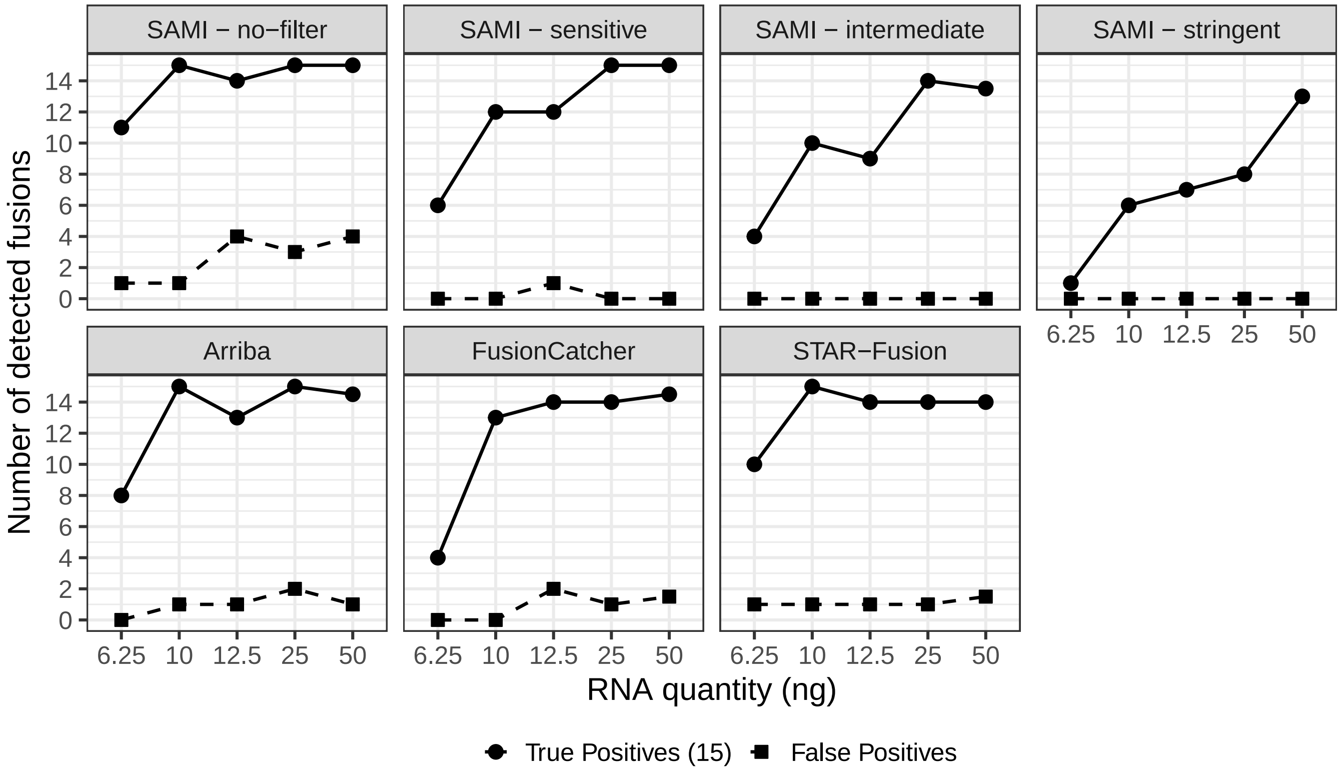

##### 2.4 Supp Figure 4: SpliceLauncher's recall without transcript list

Proportion of splicing alterations created by ASimulatoR which were detected by SAMI and SpliceLauncher, for each ASimulatoR event class and for various parameters of the two tools. Events found by a tool with slightly incorrect genomic coordinates (10 bp difference or less) were counted separately as near-matches.

In this alternative version of Figure 1d, SpliceLauncher was launched without the list of MANE transcripts used for simulating the data and had to select a transcript by itself.

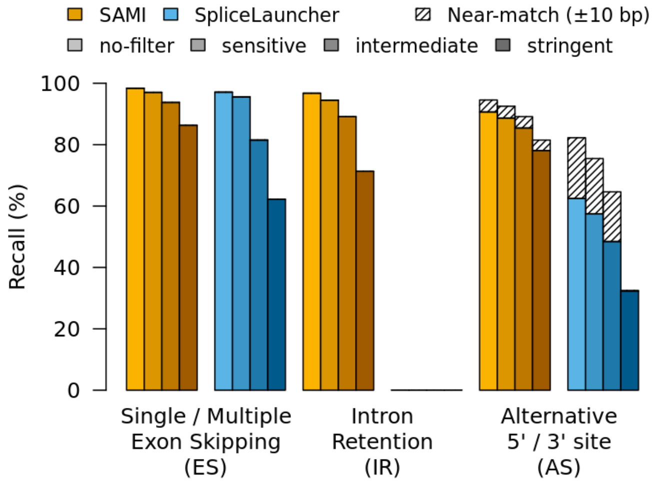

##### 2.5 Supp Figure 5: SpliceLauncher's precision without transcript list

Proportion of splicing alterations reported by SAMI and SpliceLauncher which were actually generated by ASimulatoR, split by event classes reported by each tool and for various parameters.

In this alternative version of Figure 1e, SpliceLauncher was launched without the list of MANE transcripts used for simulating the data and had to select a transcript by itself.

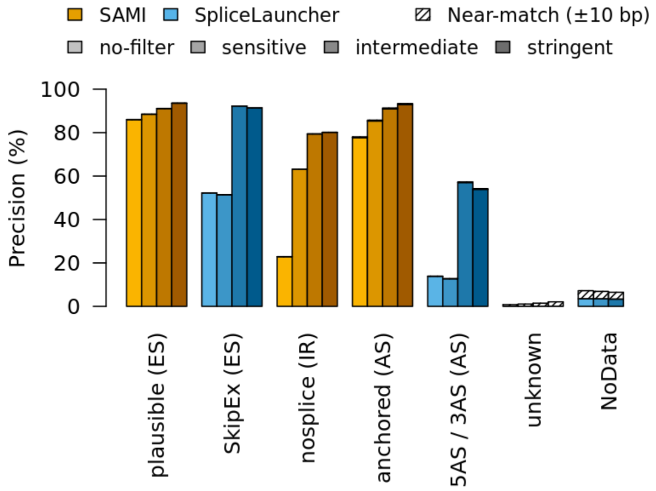

### 3 Supplemental Tables

##### 3.1 Supp Table 1 : Events expected in the Seraseq® sample

Human Genome Variation Society (HGVS) nomenclature of 17 aberrations expected in the Seraseq® commercial sample (adapted from https://www.seracare.com/globalassets/seracare-resources/pi-0710-0497-seraseq-fusion-rna-mix-v4.pdf). TMPRSS2-ERG was not considered as none of the two genes involved are captured by the "small" RNA-seq panel the sample was sequenced with.

| event | type | HGVS |
|---|---|---|
| EGFR Variant III | splicing | EGFRNM_005228.5:r.350_1150del |
| MET ex 14 Skipping | splicing | METNM_001127500.3:r.3338_3478del |
| CCDC6-RET | gene fusion | CCDC6NM_005436.5:r.1_435_RETNM_020975.6:r.2327_5617 |
| CD74-ROS1 | gene fusion | CD74NM_001025159.2:r.1_812_ROS1NM_002944.2r.5757_7368 |
| EGFR-SEPT14 | gene fusion | EGFRNM_005228.5:r.1_3207_SEPT14NM_207366.3:r.1200_3752 |
| EML4-ALK | gene fusion | EML4NM_019063.4:r.1_1763_ALKNM_004304.4:r.4125_6265 |
| ETV6-NTRK3 | gene fusion | ETV6NM_001987.4:r.1_1283_NTRK3NM_001012338.2:r.1892_3004 |
| FGFR3-BAIAP2L1 | gene fusion | FGFR3NM_000142.4:r.1_2530_BAIAP2L1NM_018842.4:r.315_3682 |
| FGFR3-TACC3 | gene fusion | FGFR3NM_000142.4:r.1_2530_TACC3NM_006342.3:r.2066_2799 |
| KIF5B-RET | gene fusion | KIF5BNM_004521.2:r.1_3231_RETNM_020975.6:r.2070_5617 |
| LMNA-NTRK1 | gene fusion | LMNANM_170707.3:r.1_762_NTRK1NM_001012331.1:r.1290_2647 |
| NCOA4-RET | gene fusion | NCOA4NM_001145260.1:r.1_1014_RETNM_020975.6:r.2327_5617 |
| PAX8-PPARG | gene fusion | PAX8NM_003466.4:r.1_1253_PPARGNM_138712.3:r.246_1892 |
| SLC34A2-ROS1 | gene fusion | SLC34A2NM_006424.2:r.1_460_ROS1NM_002944.2:r.5757_7368 |
| SLC45A3-BRAF | gene fusion | SLC45A3NM_033102.3:r.1_109_BRAFNM_004333.5:r.1206_4560 |
| TFG-NTRK1 | gene fusion | TFGNM_006070.5:r.1_851_NTRK1NM_001012331.1:r.1234_2647 |
| TPM3-NTRK1 | gene fusion | TPM3NM_153649.3:r.1_794_NTRK1NM_001012331.1:r.1234_2647 |
| TMPRSS2-ERG | off-target | TMPRSS2NM_005656.3:r.1_78_ERGNM_004449.4:r.124_5042 |

#### 3.2 Supp Table 2 : Detailed computation time

CPU and wall-clock times (in hours) measured running SAMI and SpliceLauncher (SL) on three runs from two RNA-seq panels and on ASimulatoR data.

| run | job | tool | CPU time | wall time | other | QC | UMI | splicing | STAR |
|---|---|---|---|---|---|---|---|---|---|
| small-1 | 291228 | SAMI | 15.011 | 0.507 | 0.26 | 0.756 | 1.82 | 1.496 | 10.679 |
| small-1 | 291229 | SAMI | 15.437 | 0.487 | 0.213 | 0.679 | 1.832 | 1.497 | 11.216 |
| small-1 | 291230 | SAMI | 15.251 | 0.48 | 0.219 | 0.727 | 1.75 | 1.464 | 11.091 |
| small-1 | 291231 | SAMI | 14.912 | 0.479 | 0.219 | 0.698 | 1.767 | 1.46 | 10.768 |
| small-1 | 291232 | SAMI | 14.368 | 0.469 | 0.219 | 0.696 | 1.796 | 1.46 | 10.197 |
| small-1 | 292894 | SL | 3.889 | 0.643 | 0 | 0 | 0.466 | 1.133 | 2.29 |
| small-1 | 292895 | SL | 3.053 | 0.627 | 0 | 0 | 0.429 | 1.136 | 1.488 |
| small-1 | 292896 | SL | 3.115 | 0.629 | 0 | 0 | 0.514 | 1.122 | 1.479 |
| small-1 | 292897 | SL | 3.018 | 0.62 | 0 | 0 | 0.394 | 1.122 | 1.501 |
| small-1 | 292898 | SL | 3.018 | 0.619 | 0 | 0 | 0.393 | 1.119 | 1.506 |
| small-2 | 292899 | SAMI | 12.191 | 0.36 | 0.136 | 0.437 | 1.341 | 1.042 | 9.235 |
| small-2 | 292900 | SAMI | 12.105 | 0.354 | 0.122 | 0.47 | 1.336 | 1.048 | 9.13 |
| small-2 | 292901 | SAMI | 12.103 | 0.366 | 0.129 | 0.441 | 1.357 | 1.048 | 9.129 |
| small-2 | 292902 | SAMI | 12.144 | 0.361 | 0.132 | 0.471 | 1.333 | 1.036 | 9.172 |
| small-2 | 292903 | SAMI | 12.149 | 0.354 | 0.133 | 0.437 | 1.348 | 1.043 | 9.188 |
| small-2 | 292904 | SL | 2.333 | 0.275 | 0 | 0 | 0.305 | 0.398 | 1.631 |
| small-2 | 292905 | SL | 2.056 | 0.263 | 0 | 0 | 0.298 | 0.39 | 1.368 |
| small-2 | 292906 | SL | 2.11 | 0.265 | 0 | 0 | 0.293 | 0.395 | 1.422 |
| small-2 | 292907 | SL | 2.096 | 0.261 | 0 | 0 | 0.301 | 0.387 | 1.408 |
| small-2 | 292908 | SL | 2.11 | 0.266 | 0 | 0 | 0.299 | 0.396 | 1.415 |
| small-3 | 292925 | SAMI | 14.227 | 0.406 | 0.156 | 0.598 | 1.64 | 1.078 | 10.754 |
| small-3 | 292926 | SAMI | 14.102 | 0.427 | 0.161 | 0.602 | 1.629 | 1.087 | 10.622 |
| small-3 | 292927 | SAMI | 14.521 | 0.422 | 0.16 | 0.613 | 1.61 | 1.057 | 11.081 |
| small-3 | 292928 | SAMI | 13.865 | 0.417 | 0.152 | 0.596 | 1.629 | 1.053 | 10.435 |
| small-3 | 292929 | SAMI | 14.09 | 0.419 | 0.158 | 0.578 | 1.607 | 1.053 | 10.695 |
| small-3 | 292930 | SL | 2.055 | 0.238 | 0 | 0 | 0.342 | 0.376 | 1.337 |
| small-3 | 292931 | SL | 2.044 | 0.237 | 0 | 0 | 0.338 | 0.374 | 1.332 |
| small-3 | 292932 | SL | 2.032 | 0.239 | 0 | 0 | 0.334 | 0.379 | 1.319 |
| small-3 | 292933 | SL | 2.066 | 0.237 | 0 | 0 | 0.342 | 0.373 | 1.351 |
| small-3 | 292934 | SL | 2.044 | 0.238 | 0 | 0 | 0.334 | 0.376 | 1.335 |
| large-1 | 274448 | SAMI | 89.037 | 1.971 | 4.106 | 10.534 | 26.693 | 3.837 | 43.867 |
| large-1 | 274449 | SL | 20.203 | 2.273 | 0 | 0 | 5.709 | 3.647 | 10.847 |
| large-2 | 274450 | SAMI | 100.752 | 1.996 | 4.694 | 11.921 | 30.672 | 3.992 | 49.473 |
| large-2 | 278412 | SL | 22.979 | 2.541 | 0 | 0 | 6.559 | 4.04 | 12.381 |
| large-3 | 278413 | SAMI | 104.516 | 2.343 | 5.148 | 11.626 | 29.534 | 3.981 | 54.227 |
| large-3 | 278414 | SL | 24.518 | 2.587 | 0 | 0 | 6.657 | 4.024 | 13.838 |
| ASR | 274440 | SAMI | 149.006 | 2.904 | 6.473 | 23.728 | 0 | 5.104 | 113.701 |
| ASR | 274441 | SL | 29.359 | 3.55 | 0 | 0 | 0 | 6.46 | 22.899 |
| ASR | 292951 | SL | 25.372 | 3.545 | 0 | 0 | 0 | 6.514 | 18.858 |
| ASR | 292952 | SL | 25.159 | 3.542 | 0 | 0 | 0 | 6.493 | 18.667 |
